# LeafCutter vs. MAJIQ and comparing software in the fast moving field of genomics

**DOI:** 10.1101/463927

**Authors:** Jorge Vaquero-Garcia, Scott Norton, Yoseph Barash

## Abstract

In a recent publication, Li *et al.* introduced LeafCutter, a new method for detecting and quantifying differential splicing of RNA from RNASeq data. In this work, Li *et al.* first compared LeafCutter to existing methods, then used it for a study of splicing variations and sQTL analysis from a large set of GTEx samples. While the study was elaborate and comprehensive, we want to highlight several issues with the comparative analysis performed by Li *et al.* We argue these issues created an inaccurate and misleading representation of other tools, namely MAJIQ and rMATS. More broadly, we believe the points we raise regarding the comparative analysis by Li *et al.* are representative of general issues we all, as authors, editors, and reviewers, are faced with and must address in the current times of fast paced genomics and computational research.

---

The issues we identified are all concentrated in Fig 3 of Li *et al.* [1]. We stress these do not invalidate the comprehensive GTEx analysis performed by the authors, for which they should be congratulated. These issues relate only to the comparison to other software and can be summarized by the following points:

## Usage of outdated software

Li *et al.* compared LeafCutter to three different algorithms for differential splicing analysis from RNA-Seq data: MAJIQ[2], rMATS[3] and cufflinks2[4]. Importantly, the versions of the software used were not stated. We found the versions used (MAJIQ 0.9.2, released Jan 2016, and rMATS 3.2.5) were outdated, a consequence of when the authors set up the comparative analysis for their paper submission on March 31st, 2017 (Li *et al.* personal communication, see supplementary for more details). Consequently, major software re-implementation included in rMATS 4.0 (released May 1st 2017) and MAJIQ 1.0 (released May 10th 2017), were not included in the Li *et al.* BioRxiv manuscript posted Sep 7th 2017, or the paper published on Dec 11th 2017. The usage of the outdated software packages led the authors to conclude that *“the identification of differential splicing across groups in large studies impractically slow with rMATS or MAJIQ”*. To support this claim, the authors plotted time and memory usage where both rMATS and MAJIQ were found to consume over 10GB of memory and took over 50 hours (Fig. 3b, Supplementary Fig. 3b in Li *et al.*). However, in our testing, the May 2017 versions required only a few hundred MB of memory per node, similar to LeafCutter, and while MAJIQ 1.0 was still significantly slower, rMATS was as fast as LeafCutter (see Figure 1a).

**Figure 1:**
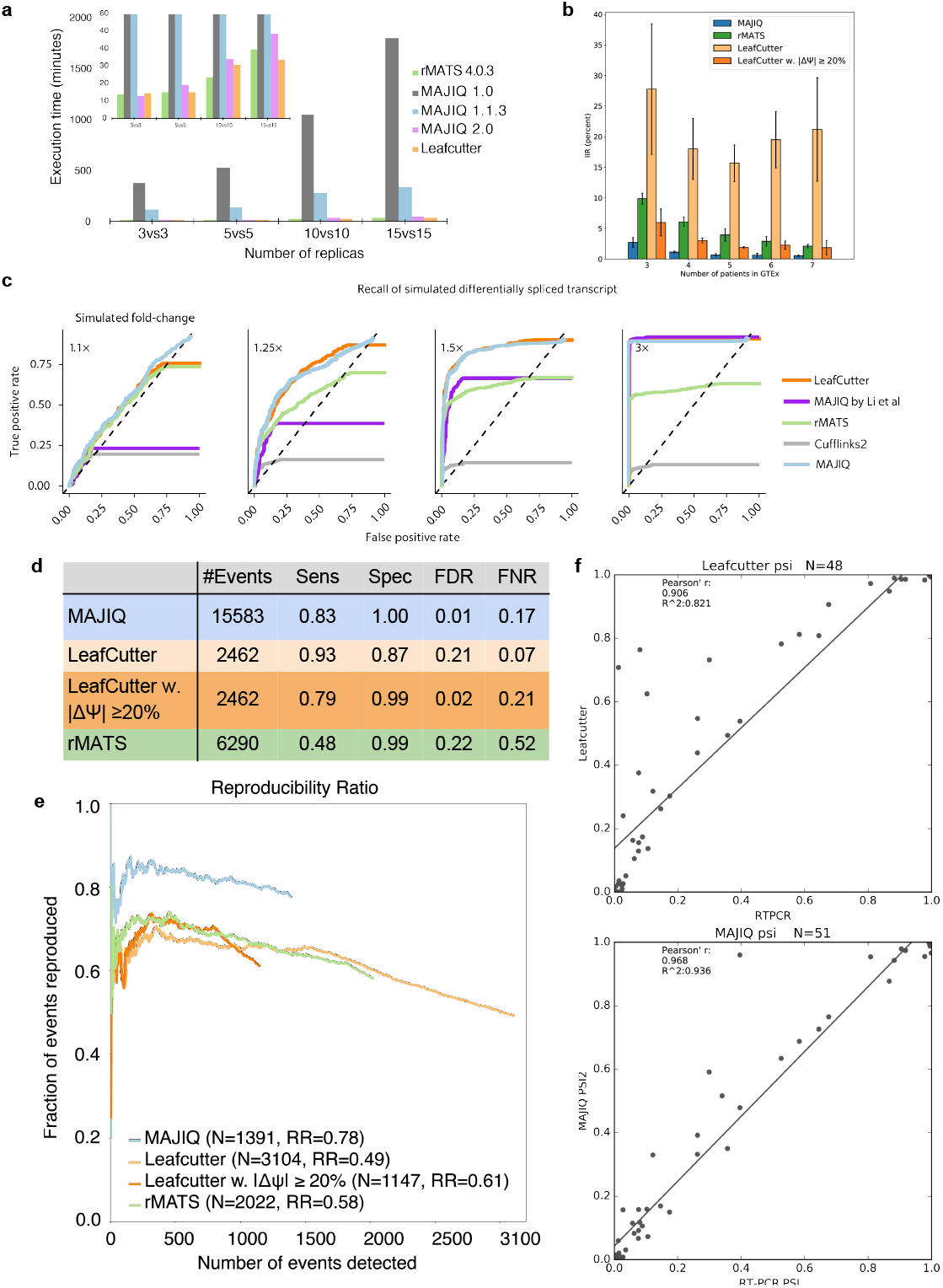
Comparative evaluation. **(a)** Running time for each algorithm, when comparing groups of different sizes. **(b)** The Intra to Inter Ratio (IIR) when using 3 7 GTEx per tissue group (skeletal muscle). The IIR, serving as a proxy for false discovery, represents the ratio between the number of differential events reported when comparing biological replicates of the same tissue (putative false positives), and the number of events reported when comparing similarly sized groups but from different conditions (here skeletal muscle and cerebellum, see main text and supplementary for details). **(c)** The original ROC plots from Li *et al.* for evaluating each method’s accuracy, with the correct execution of MAJIQ superimposed on them (blue line). The blue line was derived using scripts supplied by Li *et al.* for their data generation. **(d)** Evaluation using “realistic” synthetic datasets: each synthetic sample is created to match a real sample in terms of gene expression and a lower bound on transcriptome complexity. This simulation does involve *de-novo* events which are not captured by rMATS or intron retention (not modeled by LeafCutter). All datasets involve 3 biological replicates per group. Each method was evaluated using its own definition of alternative splicing events, so events are not directly comparable between methods. Positive events were defined as those with (|*E*[∆Ψ]| ≥ 20%), and negative events were defined as those with a small difference between the groups of (|*E*[∆Ψ]| ≤ 5%). **(e)** Reproducibility ratio (*RR*) plots for differentially spliced events between cerebellum and heart GTEx samples (*n* = 5 per group, as in Li *et al.*). The end of the line marks the point in the graph matching the number of events reported as significantly changing (*RR*(*N_A_*), see main text and supplementary). Events detected are not directly comparable as each algorithm uses a different definition for splicing events. **(f)** Evaluation of accuracy using RT-PCR experiments from [2]. Both algorithms were used to quantify Ψ using RNA-seq from [6] and RNA from matching Liver tissue was used for validation.

## Reproducibility and correct execution

In order to assess LeafCutter’s accuracy, Li *et al.* performed two main types of tests. In the first, represented by Fig 3b in the published paper, Li *et al.* assessed the distribution of p-values reported by the software. For this, the authors compared two groups of RNASeq samples which were both comprised of equal proportions of samples from two conditions (tissues). As both groups included an equal mix of the same conditions the software was expected to produce p-values that follow the null. This figure creates the false impression, also stated in the main text, that MAJIQ’s output is not well calibrated. In fact, the authors erroneously chose to use a different output type, the posterior probability for a predefined magnitude of inclusion change *C* (denoted *P* (∆Ψ *> C*)), as a proxy for p-values. This usage is wrong as p-values are derived from a null model, which is not used by MAJIQ. Instead, as evident by the shape of graph, the posterior probability produced by MAJIQ can be thought of as a soft proxy to a step function. Such an ideal step function would assign all the true ∆Ψ *> C* with probability 1 and assign probability 0 to the rest. Furthermore, when we perform such a test to assess putative false positives using GTEx samples and the setup discussed below, we find MAJIQ protected from false positives significantly better than LeafCutter (Figure 1b).

The second measure of performance employed by Li *et al.* is a receiver operating characteristic (ROC) curve, assessing the accuracy of calling differential splicing from synthetically generated RNASeq data. The authors use this analysis to conclude MAJIQ severely under-performs in any but the most extreme cases. This analysis seemed incorrect to us since the graphs show MAJIQ is not able to retrieve many events (MAJIQ’s purple line saturates quickly, see Figure 1c). As the data and scripts were not available at publication, we contacted the authors for this information and were able to repeat the analyses. We found the authors did not use MAJIQ as intended based on the users manual and analysis in [2, 5]: In order to plot such ROC curves one needs to rank events by the absolute expected inclusion change (|*E*[∆Ψ]|) and have all events reported. Reporting all events is done using the--show-all flag. Running the pipeline supplied to us by Li *et al.* and superimposing the results on the original graphs, we find there is no significant difference in this test between LeafCutter and MAJIQ (Figure 1c, blue line). However, as we detail below, in more realistic synthetic data as well as real data, we find MAJIQ outperforms both LeafCutter and rMATS by a variety of metrics.

## Realistic evaluation metrics

We found the synthetic data used to evaluate the software to be unrealistic. Among the things we noted are the use of uniform expression of isoforms, spiking a single isoform’s expression level which does not translate to any specific splicing change (∆Ψ), using a fixed number (5) of isoforms per gene in the scripts we were supplied with, usage of only a small non-random set of 200 genes, and avoiding fluctuations between individuals. The last point is especially relevant given the heterogeneous nature of the GTEx dataset. The Li *et al.* analysis also avoided any PSI or isoform specific measures of accuracy and instead assessed ROC at the gene level (spiked yes/no). In what we consider to be more realistic synthetic data mimicking biological replicates, and produced using the procedure described in [5], we find MAJIQ outperforms LeafCutter (see Figure 1d). A more complete description of the issues we found in the synthetic data can be found in the supplementary material.

A metric we found missing was reproducibility of the identified significantly changing events when biological replicates are used to repeat the analysis. Here too we found MA-JIQ’s results to be significantly more reproducible then LeafCutter’s: when using two groups of GTEx cerebellum and skeletal muscle samples, MAJIQ achieved consistently higher reproducibility irrespective of the number of events reported (Figure 1e). MAJIQ’s improved reproducibility was maintained when using biological replicates (data not shown) and when restricting LeafCutter to use a more conservative additional filter of ∆Ψ *>* 20% (compare light and dark orange lines in see Figure 1b,d). This additional filter is similar to MA-JIQ’s settings and commonly used in the RNA Biology field for defining significant splicing changes.

The evaluations in the Li *et al.* analysis also did not include any experimental validation or accuracy measure by RT-PCR. While RT-PCR can suffer from biases as well, careful execution in triplicates is considered the golden standard in the RNA field. To make those accessible, we and others have made datasets of such experiments readily available online [2, 5]. Using those datasets, we found LeafCutter to be significantly less accurate when compared to MAJIQ (*R*^2^ 0.821 vs 0.936 in Figure 1f) and when compared to rMATS. Moreover, we believe these results highlight an inherent issue in LeafCutters output: while useful for sQTL detection (the use case for which LeafCutter was originally designed for), LeafCutters cluster of introns are a mathematical construction. The clusters do not correspond directly to a biological entity or to ratios of isoforms which necessarily add up to one. As such, it is not clear how LeafCutter’s output should be translated to actual inclusion values, or be used for primer design.

Finally, there are several important elements of LeafCutter which were not discussed in the Li *et al.* comparative analysis. Those include the limited granularity of intron clusters, which can cover large portions of genes (see number of events reported in Figure 1d as a crude indication of this issue), and the fact that intron retention (IR) is not modeled by LeafCutter. The latter has been shown to be of particular importance in the brain, a focus of Li *et al.* s analysis. For example, in a preliminary analysis of brain GTEx samples we found almost 10% of the differentially spliced events to involve differential IR (data not shown).

## Conclusions

In conclusion, our evaluation supports the Li *et al.* assertion that LeafCutter is an efficient method for differential splicing and particularly sQTL analysis, for which it was originally constructed. We note that our analysis does not imply purposeful misrepresentation of competing software (see time-line in supplementary), nor does it invalidate the comprehensive analysis of GTEx samples performed by the authors. Nonetheless, we conclude that Li *et al.* misrepresented other software and their relative performance, namely rMATS and MAJIQ. This misrepresentation was a result of using outdated software, lack of proper documentation (software versions, scripts), incorrect usage, and what we view as lacking evaluation criteria.

Nature Research journals have been strong advocates for reproducible science (*cf* [7, 8]. We believe that adherence to relevant reproducibility guidelines could have helped prevent many of the issues we identified. But beyond the need to follow reproducibility guidelines, other important questions arise. For example, what evaluation criteria should we use? Which software version should we run? What happens if a competing software is updated during a review? Are we obliged to change the manuscript then? Are we also obliged to address pre-prints? Should reviewers be allowed to use pre-prints to scrutinize a manuscript or a grant application? Can authors use pre-prints to base their scientific claims in other papers? And what is the role of the editor in such cases? We found ourselves struggling with those questions ourselves. For example, we faced reviewers of grants and papers which scrutinized our and collaborators work based on this inaccurate representation of MAJIQ, even when it was only a pre-print. We published improvements to the MAJIQ algorithm, which was released as a pre-print back in January 2017, but we choose not to discuss algorithmic enhancements here, as it was only formally published on December 2017. As another example, during the preparation of our Norton *et al.* manuscript [5], competing methods released updates. We decided to delay submission, rerun all comparative analyses, and consequently removed specific claims. How should such software changes be handled by reviewers and editors? As it stands, we are painfully aware that MAJIQ itself represents a moving target as it is actively being developed. For example, the current version (1.1, released March 1st 2018) is much faster than the previous 1.0 release, and the new version (MAJIQ 2.0, manuscript in preparation) compares well in running time to LeafCutter while still retaining MAJIQs added features (see Figure 1a). Consequently, we found even more recently published splicing analysis methods such as SUPPA2[9] and Whippet[10] also used outdated MAJIQ software (ver 0.9, released Feb 2016) or did not report versions. Such situations of constantly evolving software is not uncommon in genomics, with methods such as rMATS[11, 3], Salmon[12], EdgeR[13], and DESeq[14, 15] serving as good examples. In such settings, the decision on which version to use (and documenting it) can critically affect results.

Finally, we want to point out an important take home message for our community. In this fast moving field of genomics, with evolving standards and procedures, mistakes are bound to happen even as we strive to minimize them. The question, then, is how do we handle mistakes? In preparing this response, we found the online discussion around the MAJIQ vs LeafCutter analysis, including personal communications with the authors, to all be helpful and constructive. The authors responded to our queries and supplied their scripts; other researchers even suggested to use this case to write a guide for software comparisons. We believe this sort of constructive response sets an excellent example, and hope we as a community will keep this positive approach in the future for the benefit of all.

## Addendum

- We would like to thank Li *et al.* for their email correspondence, supplying their original scripts, providing feedback to this manuscript and sending us the time-line included in the supplementary information.
- All scripts and data used to generate the results presented here are available at: https://bitbucket.org/biociphers/vaquero_norton_2018/
- Competing Interests: The authors declare that they have no competing financial interests.

## Supplementary Information

### Tool use

All the tools used were the last version found by the authors on August 2018, specifically rMATS 4.0.3 and Leafcutter v0.2.7-18-g7ad3ee1 (hash:7ad3ee1eba7a9bf6783f24b4a5b4b8f6aac91c9c). Running times reported are based on a 32-core Centos 7 machine with 64 GB.

All the runs are done using each software provided options for multiprocessing/multithreading set to 16. This is achieved using rMATS options *–nthreads* and *–tstats*; Leafcutter option *-p*; and MAJIQ *–nthreads* or *-j*. Figure 1a reports execution time, calculated as time from start with bam files to producing each tool’s output. This means human readable text files for the three tools, with default parameters except for the threading options explained before. We also made the following simplifying assumptions in favor of the tools we compared against. First, we assumed an ideal execution framework for leafcutter, where a user was able to perfectly parallelize the supplied bam2junc.sh, which does not include parallelization, into parallel execution of 16 files without any computational overhead, *i.e.*, 1*/*16th time of a single thread execution. We also note that the amount of information provided by each tools is different since Leafcutter doesn’t quantify intron retention and rMATS *denovo* detection is limited to known introns and exons. Still, we did not remove de-novo from MAJIQ and ran it with regular settings.

All the execution scripts used on this work will be available publicly upon publication.

### Software and manuscript time-line

The following time line, supplied to us by Li *et al.*, documents the times in which software and manuscripts were posted:

- Feb 1, 2016: MAJIQ paper comes out in eLife, MAJIQ version 0.9.1.
- March 16, 2016: First LeafCutter manuscript posted on bioRxiv. It does not include accuracy analysis compared to other software.
- April 13, 2016: LeafCutter manuscript rejected without review.
- April 29, 2016: Large scale LeafCutter based sQTL analysis published [16].
- March 31, 2017: LeafCutter manuscript submitted to Nature Genetics including accuracy analysis compared to MAJIQ and rMATS.
- May 1, 2017: rMATS-turbo, is released as a Docker installation.
- May 10, 2017: MAJIQ 1.0 released.
- September 7, 2017: Updated LeafCutter manuscript posted on bioRxiv, including comparative analysis.
- November 15, 2017: rMATS-turbo released as v.4.0.1 (non Docker installation).
- December 11, 2017: Blog post, with an initial examination of potential issues with the comparative analyses in the LeafCutter’s BioRxiv preprint, is posted [17].
- December 11, 2017: LeafCutter manuscript published in Nature Genetics.
- May 17, 2018: MAJIQ 1.1.3 released.

### Synthetic data

We found several aspects of the synthetic data used in Li *et al.* to be unrealistic or limiting. Specifically:

1. Gene set: The set of genes selected for the analysis was small (200) and nonrandom (the first on chromosome 1).
2. Gene isoform number and consequent transcriptome complexity: In the scripts we were given by the authors each gene got exactly 5 isoforms.
3. Gene expression levels: The expression levels were not made to mimic those observed in real data (GTEx).
4. Relative expression levels of isoforms: The authors chose to set all isoforms to the same expression level, except a single isoform which was “spiked” at varying expression folds (1.1, 1.25, 1.5, 3, 5). In real data, however, the distribution across isoforms is highly skewed [18], resulting in typically few dominant isoforms. Consequently, the observed inclusion level, or Ψ, in binary events has a typical “U” shape (see for example, supplementary figure 1 in [11] for a good illustration of that across 16 human tissues). MAJIQ utilizes this as a prior over Ψ as part of its model.
5. Spiked expression changes do not translate directly to splicing changes: Both MAJIQ and rMATS have been designed to detect changes in splicing at a predefined level (*e.g.*, 20% change, or ΔΨ = 0.2). Thus, one would expect their evaluation in an ROC to be based on positive and negative cases that meet that criteria. However, the testing shown in Figure 1c is not based on such labels and therefore does not capture true sensitivity and specificity for these algorithms. Furthermore, there is no clear way to translate spiked expression to changes in inclusion levels. For example, let us consider a change of 10% in the expression of one of the isoforms. In the best case scenario, an isoform whose expression is changed is solely responsible for the inclusion of a specific junction (we note this is not guaranteed by the procedure employed by Li *et al.*). In such a case, assuming the expression of all other isoforms amounts to *Y*, then 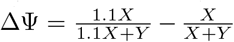. Thus, in the best case scenario under Li *et al.* setup with 1.1*X* expression spiking and using 5 equally expressed isoforms we get ∆Ψ ~ 0.016. If the distribution of isoforms across the junctions is different, this can be easily smaller. It is true one can still rank events by their estimated ∆Ψ (this was not done for MAJIQ) but we hold the test is still not reflective of the positive or negative labels by ∆Ψ which MAJIQ and rMATS are designed to capture, *i.e.*, quantitative changes in splicing.
6. Limited fluctuations of inclusion between replicates and non replicates: In real data, the observed inclusion levels are expected to naturally vary between individuals. This is true for biological replicates and even more so for heterogeneous data such as GTEx where a set of tissue samples come from different donors. Figure 3) illustrates the fluctuations of inclusions levels in biological replicates (blue), GTEx samples (green), and Li *et al.* synthetic data (red).

The synthetic data we employed, originally used in [5], was developed to mimic biological replicates (Figure 3, orange). In this data, the expression levels and transcriptome variations for over 3000 genes were simulated (Figure 3, orange) to match real samples from [19] (Figure 3, blue). We used the most complex splicing event we can find for each gene based on reliably-detected (multiple reads in multiple positions) junction-spanning reads by STAR [20]. This enabled us to define a lower bound on each gene transcriptome complexity. The expression level of each isoform to be simulated was set by the genes overall FPKM and the raw junction-spanning read ratios in the real sample to avoid biasing estimations towards any specific algorithm. This allowed us to simulate not just cassette exons or binary events but also complex variations, which both MAJIQ and LeafCutter are geared to capture. Simulated data was generated by our colleague Dr. Greg Grant using BEERS [21]. More details can be found in [5].

**Figure 2:**
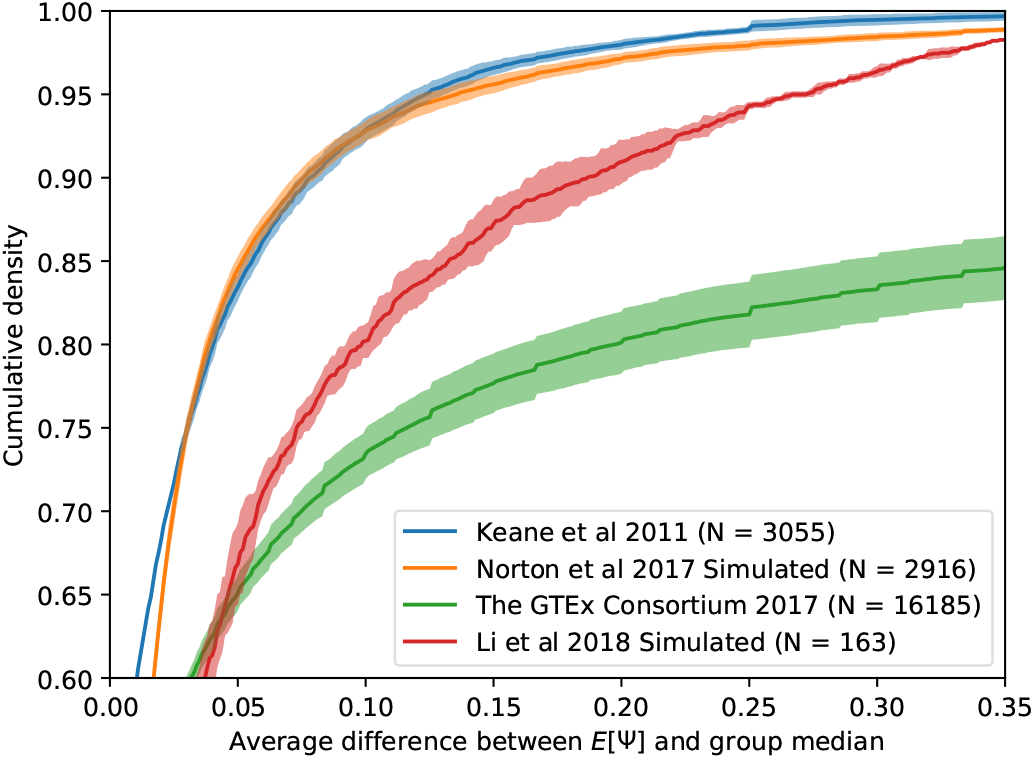
Local Splicing Variations (LSV). Exons are blue, splicing events are black edges. The LSV formulation we developed captures the classical, binary, AS events involving two alternative junctions (top) but also more complex (non binary) splicing variations and effects of alternative gene start/end (bottom).

### Real data

#### Reproducibility plots (RR)

RR plots as those shown in Figure 1f follow a similar procedure to that of irreproducible discovery rate (IDR) plots[22], used extensively to evaluate ChIP-seq peak callers [23]. Briefly, we ask the following simple question: Given an algorithm *A* and a dataset *D*, if you rank all the events that algorithm *A* identifies as differentially spliced 1 &#2026; *N_A_*, how many would be reproduced if you repeat this with dataset *D*, comprised of similar experiments using biological/technical replicates? The *RR*(*n*) plot, as shown in Figure 1e, is the fraction of those events that are reproduced (*y*-axis) as a function of *n ≤ N_A_* (*x*-axis), with the overall reproducibility of differentially spliced events expressed as *RR*(*N_A_*) (far right point of each graph in Figure 1f). Briefly, for rMATS and LeafCutter events reported as differentially spliced were ranked by their associated *p*-value with a cut-off of 0.05 as in Li *et al.* and recommended by authors. We also tested adding an additional filtering step of |[∆Ψ] ≥ 0.2|. to make selection criteria closer to what is applied by default to MAJIQ (see main text). For MAJIQ, differentially spliced events were filtered for high confidence using a posterior threshold *P* (|∆Ψ| ≥ 0.2) ≥ 0.95 and ranked by |*E*[∆Ψ] ≥ 0.2|. A more detailed description of *RR* can be found in [2, 5].

**Figure 3:**
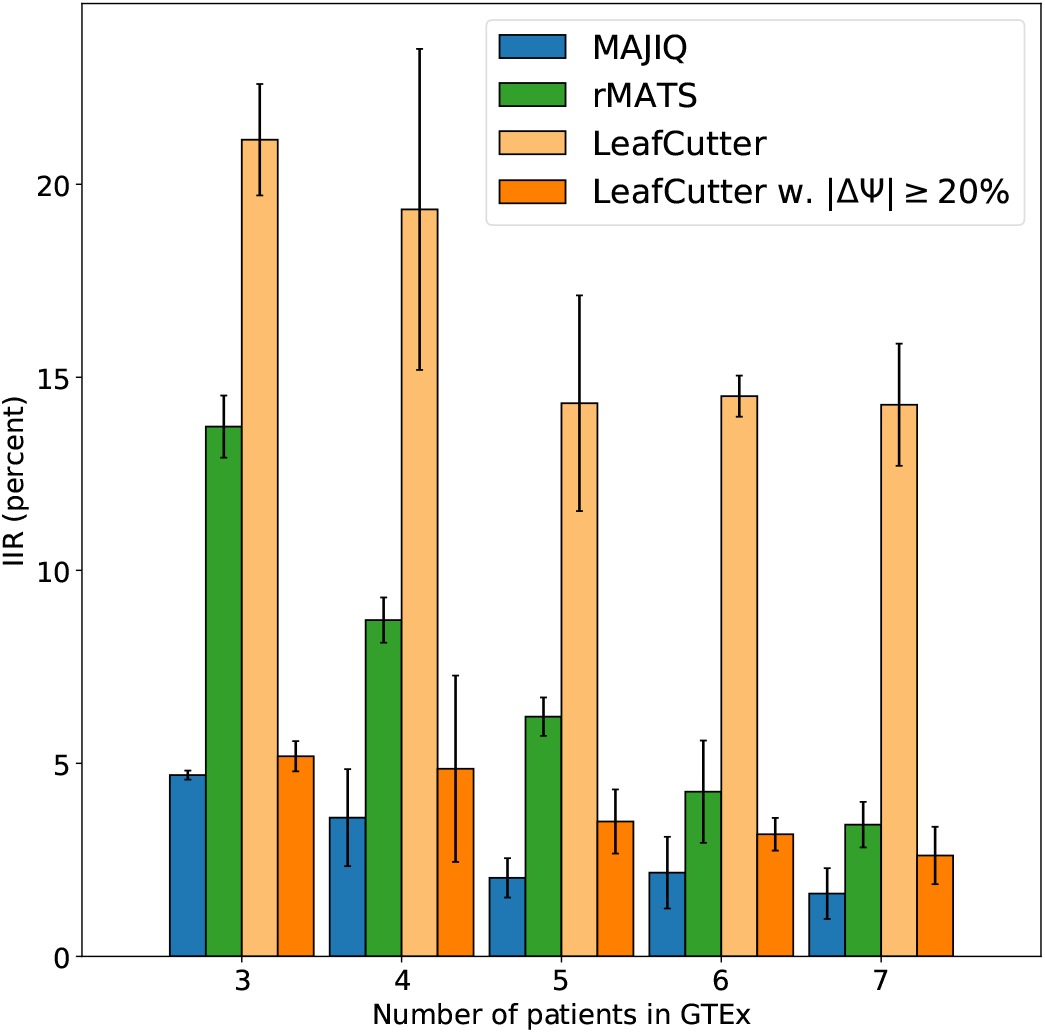
IIR for GTEx samples from cerebellum. Analysis is equivalent to that used for skeletal muscle samples in Figure 1

#### Intra to Inter Ratio (IIR)

Reproducibility alone is not sufficient to establish accuracy. For example, an algorithm can be extremely reproducible but highly biased. To get a better sense of possible levels of false positives, we devised the following test: We compare similarly-sized groups of the same condition (e.g. brain vs. brain or liver vs. liver) and compute the ratio between the number of events identified as significantly changing in such a setting (*N_PFP_*) to the number of events identified between the groups of different conditions (*N_A_*, e.g. brain vs. liver). We term this the intra to inter ratio 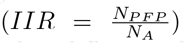, as intuitively it computes the ratio between the number of events identified as different when comparing groups that share the same label or condition (intra) and similarly sized groups of different conditions (inter). We note the intra set are not necessarily false positives, but we consider them as putative false positives (PFP) with respect to the change we are interested in between conditions. More details can be found in [5]. Notably, the IIR test is similar to the one used by Li *et al.* to assess false positives (Fig3b in [1]). In their test setting, aiming to assess FDR, Li *et al.* mix the samples such that the condition of interest are balanced between the two groups (*e.g.*, 3 liver and 3 brain samples vs another set of 3 liver and 3 brains).

**Figure 4:**
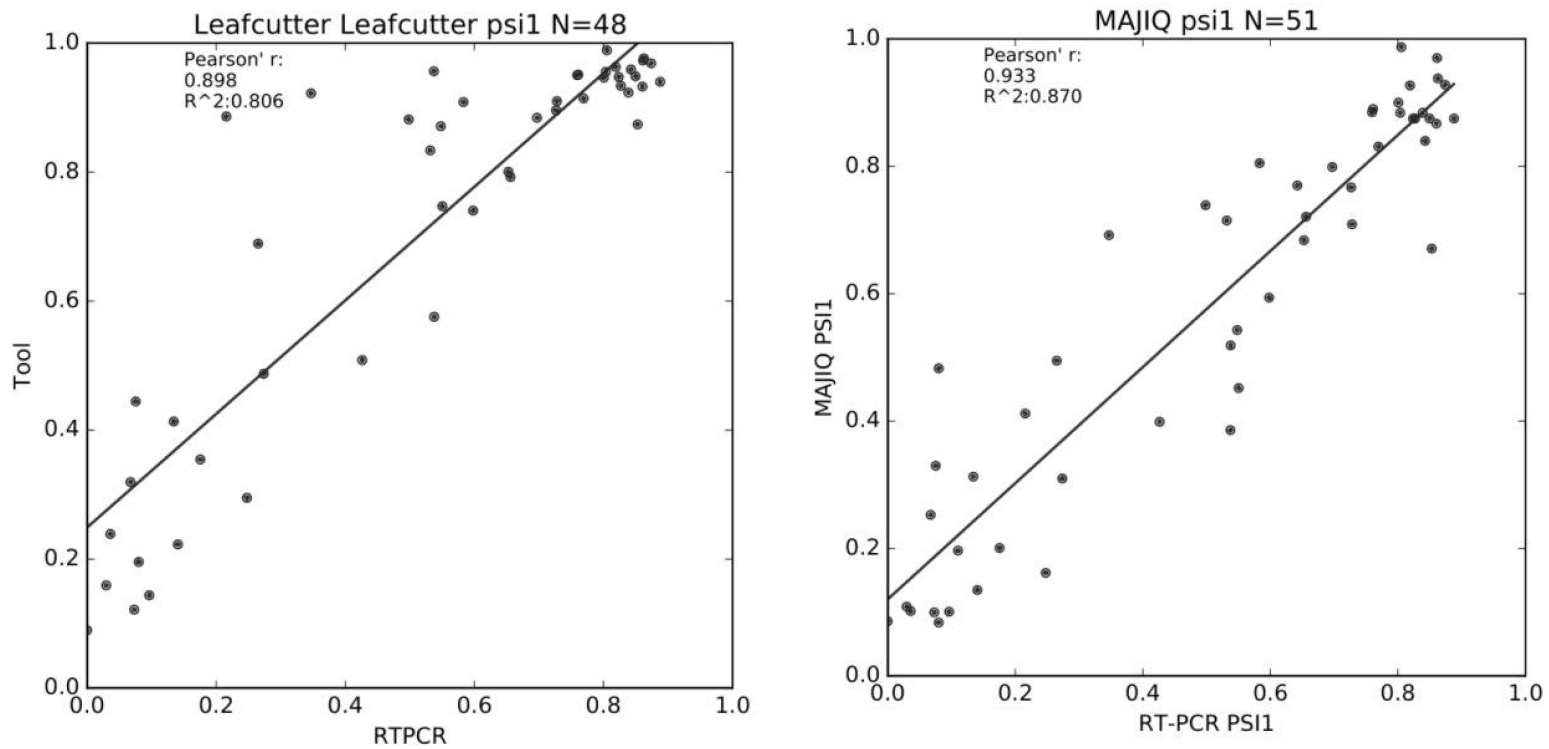
Additional evaluation using RT-PCR. Here RT-PCR experiments from [2] were used to assess LeafCutter (left) and MAJIQ (right) inclusion level (Ψ) quantification. Both algorithms were used to quantify Ψ using RNA-seq from [6] and RNA from matching Cerebellum tissue was used for validation. The set of events shown here were selected to be binary (involving only two alternative junctions), known (both junctions annotated in the transcriptome), and differentially spliced between mouse cerebellum and liver (see [2] for details).

### RT-PCR

While the above tests are important and informative, these are focused on detecting splicing changes (∆Ψ) and do not address the question of how accurate the actual inclusion levels, or Ψ, quantified by the algorithm. This is particularly relevant for the analysis in Li *et al.* for sQTL, as this is done using Ψ in each sample and not ∆Ψ between groups. Moreover, the manuscript did not include experimental validations or independent measure of accuracy. For this, RT-PCR experiments are considered the gold standard in the RNA field. We note that RT-PCR experiments are not free of potential pitfalls and biases. For example, many available RT-PCR from publications may include quantitative values for Ψ but were only designed to be used qualitatively (i.e. change vs no change). The events selected for RT-PCR validation can be highly biased (e.g. only cassette exons), and naturally these are limited throughput experiments. Nonetheless, we assert that including carefully executed RT-PCR is important for methods that make strong claims about splicing quantification, and such datasets have been made available by us and others [2, 5] - see Figure 1g and Figure 4 for comparative analysis of LeafCutter, MAJIQ, and rMATS based on those. The events shown here were pre-selected to be classical cassette events which exhibit a clear change between tissues to enable comparison of MAJIQ and rMATS in [2].

